# The vagus nerve mediates the suppressing effects of peripherally administered oxytocin on methamphetamine self-administration and seeking in rats

**DOI:** 10.1101/2019.12.18.880914

**Authors:** Nicholas A. Everett, Anita J Turner, Priscila A Costa, Sarah J. Baracz, Jennifer L. Cornish

**Affiliations:** Centre for Emotional Health, Department of Psychology, Macquarie University, NSW, Australia

## Abstract

**Background:** The neuropeptide oxytocin has emerged as a promising pharmacotherapy for methamphetamine (METH) addiction, and clinical trials of intranasal oxytocin are underway. However, there is debate as to how peripherally administered oxytocin alters brain signaling to modulate addiction processes. Interestingly, there is evidence for functional interactions between peripheral oxytocin administration and the vagus nerve. Therefore, this study investigated whether the effects of peripherally administered oxytocin require vagal signaling to reduce METH self-administration and reinstatement of METH-seeking behaviours.

**Methods:** Male and female Sprague-Dawley rats underwent surgery for jugular catheterization and either subdiaphragmatic vagotomy (SDV) or a sham operation. Rats were trained to self-administer METH, and the effect of peripherally administered oxytocin on METH intake was assessed. Rats then underwent extinction, and effects of oxytocin were assessed on cue- and METH-induced reinstatement of METH-seeking.

**Results:** Oxytocin treatment robustly attenuated METH intake in both sexes. Strikingly, SDV entirely prevented the suppressant effect of oxytocin (0.3 mg/kg) on METH intake, and partially prevented the effects of 1 mg/kg oxytocin in both sexes. After extinction, SDV impaired the suppressing effects of oxytocin on cue- and METH-primed reinstatement in males, but not females. SDV was functionally confirmed by measuring food intake following administration of the vagal dependent peptide, cholecyostokin-8.

**Conclusion:** Our data suggest that vagus nerve signaling is required for the anti-addiction effects of peripherally administered oxytocin, and that this vagal dependency is partially mediated by sex and drug withdrawal. This study has considerable implications for the applicability of oxytocin as a therapy for METH use disorder for both sexes.

## Introduction

Abuse of the addictive psychostimulant methamphetamine (METH) is associated with numerous psychosocial and physical risks^1^. Current psychosocial interventions for METH use disorder are not widely available and are minimally effective^2^, and despite significant advances in understanding the neurobiology of METH addiction, no effective pharmacotherapies have been developed^3,4^. As the pharmacological agents based on these advances progress through clinical trials and continue to produce largely negative findings^5^, there is an urgent need to develop new pharmacotherapies.

The neuropeptide oxytocin is a promising novel therapy for METH addiction^6–10^. Preclinical studies have shown that peripherally administered oxytocin can suppress METH self-administration^11^, and relapse-like behavior elicited by METH-cues^12–14^ or by METH re-exposure^15–17^. When administered chronically during abstinence from METH, oxytocin also protects against cue- and METH-induced relapse in male and female rats^18^. Clinical research also indicates efficacy of intranasally delivered oxytocin in reducing alcohol and marijuana cravings^19–21^. A synthesis of these preclinical and clinical findings has laid the groundwork for the first clinical trial using intranasal oxytocin in METH-dependent users^22^ (NCT02881177). However, intranasal oxytocin treatment has not always been effective (see ^23^ for review), which raises concerns as to whether intranasal administration in humans will recapitulate the effects of peripheral administration in rodents.

Peripheral administration of oxytocin evokes powerful anti-addiction effects, however, it is unclear how the large peptide accesses the brain following peripheral dosing to modulate neural circuits which control addiction processes^24^. Indeed, as little as 0.002-0.005% of subcutaneously or intranasally administered oxytocin can be measured in the brain^23,25^, while supraphysiologic concentrations of circulating oxytocin are measured following treatment with these peripheral doses^26^. As oxytocin receptors are expressed widely throughout the body^26^, it is possible that peripherally administered oxytocin may signal to the brain via a peripherally mediated mechanism.

The vagus nerve bidirectionally conveys information between the brain and the peripheral organs, with up to 80% of afferent fibers signaling to the brain^27^. The primary recipient of these vagal afferents is the nucleus of the solitary tract (NTS), which has noradrenergic projections to various brain regions involved in addiction^28^, as well as projections to the paraventricular (PVN) and supraoptic (SON) nuclei of the hypothalamus which appose oxytocin-secreting neurons^29^. This ascending vagal pathway is therefore well positioned to regulate the function of the endogenous oxytocin system, as stimulation of vagal afferents elevates circulating oxytocin levels^30^ and increases neuronal activity in the NTS and PVN^31^. Importantly, abdominal vagal fibers express oxytocin receptors^32^, as well as vasopressin V1A receptors^33^ which oxytocin also acts on to reduce addictive behaviours^15^, and are excited by oxytocin in *ex vivo* preparations^32^. Together, these studies provide a framework in which the vagus nerve is a gateway for peripherally administered oxytocin to signal to the brain to regulate behaviour.

Stimulation of the vagus nerve is being investigated as a therapy for a variety of psychiatric conditions^34^ which oxytocin also has efficacy for treating, including depression^35,36^, anxiety^37,38^, and post-traumatic stress disorder^39,40^. Further, vagus nerve stimulation (VNS) has preclinical efficacy for reducing addiction-like behaviours. Specifically, chronic VNS during daily extinction sessions from heroin or cocaine self-administration reduced relapse-like behaviours, and modulated activity in cortico-limbic structures involved in addiction^41,42^. Collectively, these studies indicate that peripherally administered oxytocin can stimulate the vagus nerve, and that VNS is able to modulate neural pathways involved in psychiatric conditions which oxytocin may also ameliorate. However, it is unknown whether the anti-addiction effects of peripherally administered oxytocin *require* vagal signaling. Therefore, elucidation of the relationship between exogenous oxytocin, vagal signaling, and addiction is an important step towards effective clinical translation of oxytocin as a pharmacotherapy.

Here we investigated whether the inhibitory effects of peripherally administered oxytocin on METH addiction require vagal signaling, by surgically resecting the subdiaphragmatic vagus nerve prior to volitional METH intake, reinstatement, and oxytocin treatment. Further, as preclinical evidence suggests that female rats display higher METH addiction behaviours^43^, and as the efficacy of oxytocin for reducing METH addiction may be more potent in females^44^, we investigated this hypothesized interaction of oxytocin and the vagus nerve in both sexes.

## METHODS AND MATERIALS

### Subjects

Sprague Dawley rats (*n*=20/sex) were housed in groups of 3-4 under a 12h light-cycle (lights on 08:00) in open top cages (64×40×22cm) with paper bedding, plastic tunnel, and wooden sticks. Standard chow and water were available *ad libitum* until experimental procedures began. Procedures were approved by the Macquarie University Animal Ethics Committee.

### Liquid Diet

Two days prior to surgery, solid rat chow was removed from home cages, and was replaced with a nutritionally complete liquid diet (Teklad Diet 09215) made in 90% home-cage water and 10% sweetened condensed milk for palatability. This is essential for the survival of vagotomised animals. Our observations are that subdiaphragmatic vagotomy (SDV) results in extreme gastric stasis, stomach distension, and precipitous weight loss without a liquid diet.

### Subdiaphragmatic vagotomy and intravenous catheter implantation surgeries

Rats were anaesthestised with isoflurane and were randomly assigned to receive either sham surgery, or SDV. Sham rats had their bilateral subdiaphragmatic vagi isolated but not damaged, while SDV rats had a section of each nerve from the diaphragm to the stomach cut^45^. All rats were then implanted with a jugular vein catheter externalized to their midscapular region. Rats recovered for 7-10 days, and their catheters were flushed daily with heparinized cephazolin in saline. Detailed surgical procedures are described in the Supplement.

### Methamphetamine Self-administration Training

Rats were trained to self-administer METH (0.1 mg/kg/0.05 ml infusion) during 14 daily fixed ratio 1 (FR1) sessions (2h/day). Self-administration chambers were equipped with infrared beams for locomotor tracking, two illuminated nose-pokes (“active” and “inactive”), a house-light, and infusion pump. Active nose-pokes triggered the infusion pump and illuminated the nose-poke for 3 seconds. A 20-second time-out was signaled by dimming of the house-light. Inactive nose-pokes had no consequences.

### Tests for the interaction of SDV and oxytocin on methamphetamine self-administration

The effect of oxytocin (0.3, 1.0 mg/kg; ChinaPeptide) or vehicle (0.9% saline) was tested on METH self-administration. Treatments were delivered in the home cage via intraperitoneal (IP) injection 30 minutes prior to the session. Treatments were counterbalanced across 4 test days, according to a Latin square design, with at least two self-administration days between each test.

### Extinction

Rats then underwent behavioural extinction sessions for 14 days (1h/day), during which nose-pokes had no consequences. Rats received IP saline immediately prior to the final two extinction sessions.

### Tests for the interaction of SDV and oxytocin on cue-induced and methamphetamine-primed reinstatement

Following extinction, rats underwent two cue-induced reinstatement tests in the presence of methamphetamine-cues. The 1h sessions began with a single non-contingent cue presentation (30 seconds into session). Thereafter, active nose-pokes produced contingent presentation of the methamphetamine-cue. Thirty minutes prior to these tests, rats received saline or oxytocin (0.3 mg/kg IP). As repeated cue-induced reinstatement testing would result in extinction of the operant behaviour, only a single oxytocin dose was tested compared with vehicle. The lower dose was chosen due to its vagal-dependent effects on METH self-administration. Rats then underwent three METH-primed reinstatement tests, in which rats received saline or oxytocin (0.3, 1.0 mg/kg IP) 30 minutes prior to a METH injection (1 mg/kg IP), and then were placed immediately into the chamber under extinction conditions. Each reinstatement test was separated by 2-3 extinction days. Rats did not proceed to the next test day until they reached extinction criteria (<10 active pokes). Treatments were counterbalanced within each reinstatement type.

### Functional test of vagotomy

For the week following reinstatement tests, food was removed from home cages (16h deprivation from 10pm–2pm^46^), and rats were trained to consume their food in a metabolic chamber (1h/day at 2-3pm). Food was returned to their home cage for another 7 hours. Once consumption was stable across 2 days, rats were tested for their intake following treatment with saline or CCK-8 (4μg/kg IP; Abcam, ab120209) 30 minutes prior to being placed in the chamber. Food consumed after 1h was recorded, and effects of CCK-8 on intake is expressed in text as percent reduction of intake compared to the day prior (percentage reduction of intake = −100(intake day prior – intake following CCK-8)/intake day prior).

### Statistical Analysis

We used factorial analyses of variance and *t* tests using SPSS (Version 25, General Linear Model Procedure). Significant main and interaction effects (*p*<0.05) were followed by post-hoc tests (Fisher’s least significant difference). Significant effects crucial for data interpretation are indicated in figures, with the remainder in Supplemental Results. For self-administration data, the analysis was performed on METH infusions earned. For reinstatement data, analysis was performed on active nose pokes. The first 30-minutes of sessions were analysed, as the effects of oxytocin dissipate after this time (Supplemental Figure 2).

## RESULTS

### Excluded Animals

One female sham rat lost catheter patency during self-administration, so was excluded. During extinction, one female sham rat was euthanized due to poor health. Final sample sizes were *n*=10 for male sham, male SDV, and female SDV, and *n*=8 for female sham.

### Acquisition of methamphetamine self-administration

All rats acquired METH self-administration (>10 infusions/session; Figure 1). Total METH intake did not differ based on a sex×surgery interaction (F(1,34)=0.024, *p*=0.899), sex (F(1,34)=0.059, *p*=0.811) or surgery (F(1,34)=1.005, *p*=0.271).

**Figure 1.**
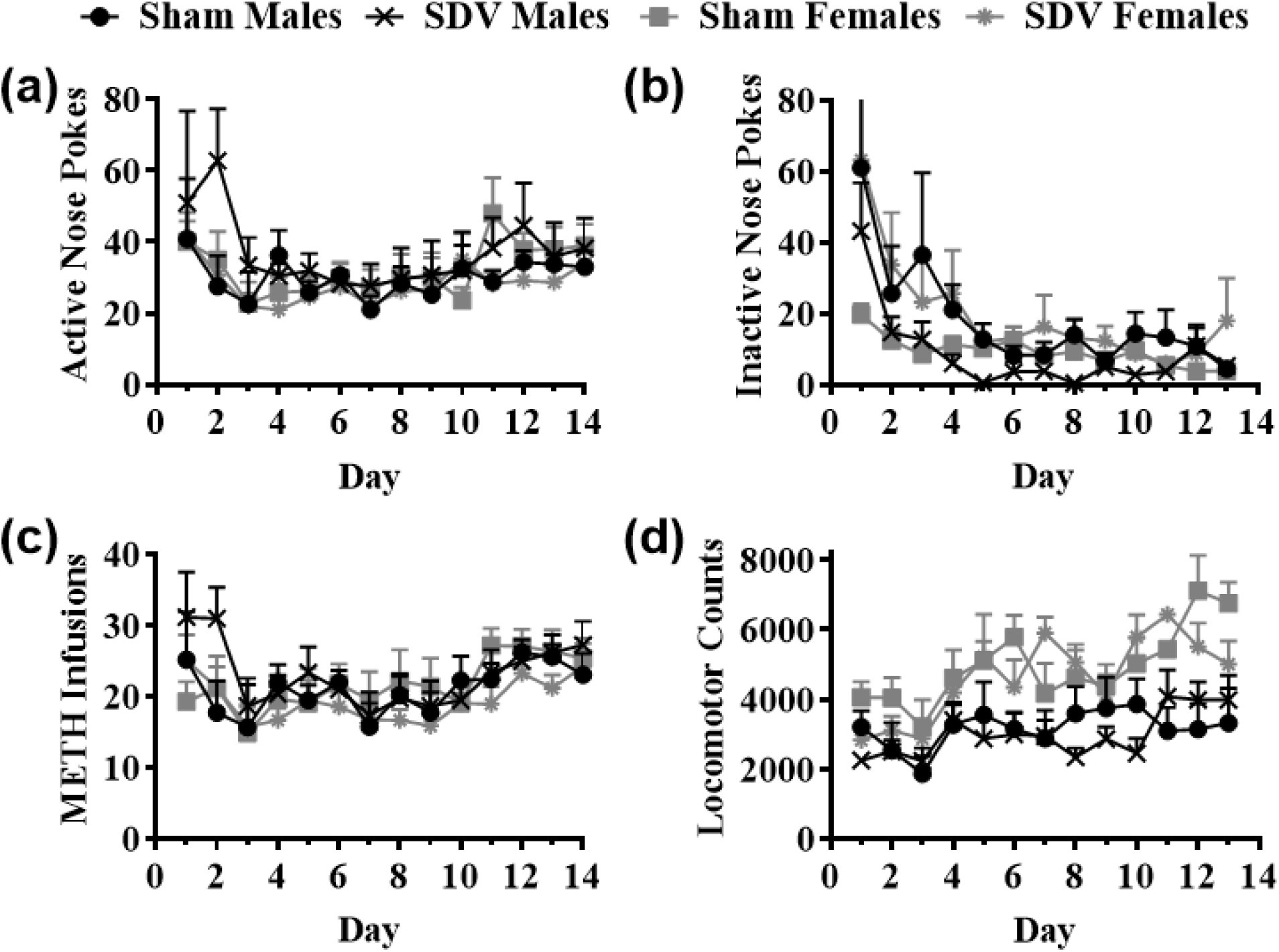
Acquisition of METH self-administration. (a) active nose pokes; (b) inactive nose pokes; (c) METH infusions; (d) locomotor counts over 14 consecutive days of intravenous METH self-administration between sham operated and vagotomised (SDV) male and female rats. Data are mean ± SEM.

**Figure 2.**
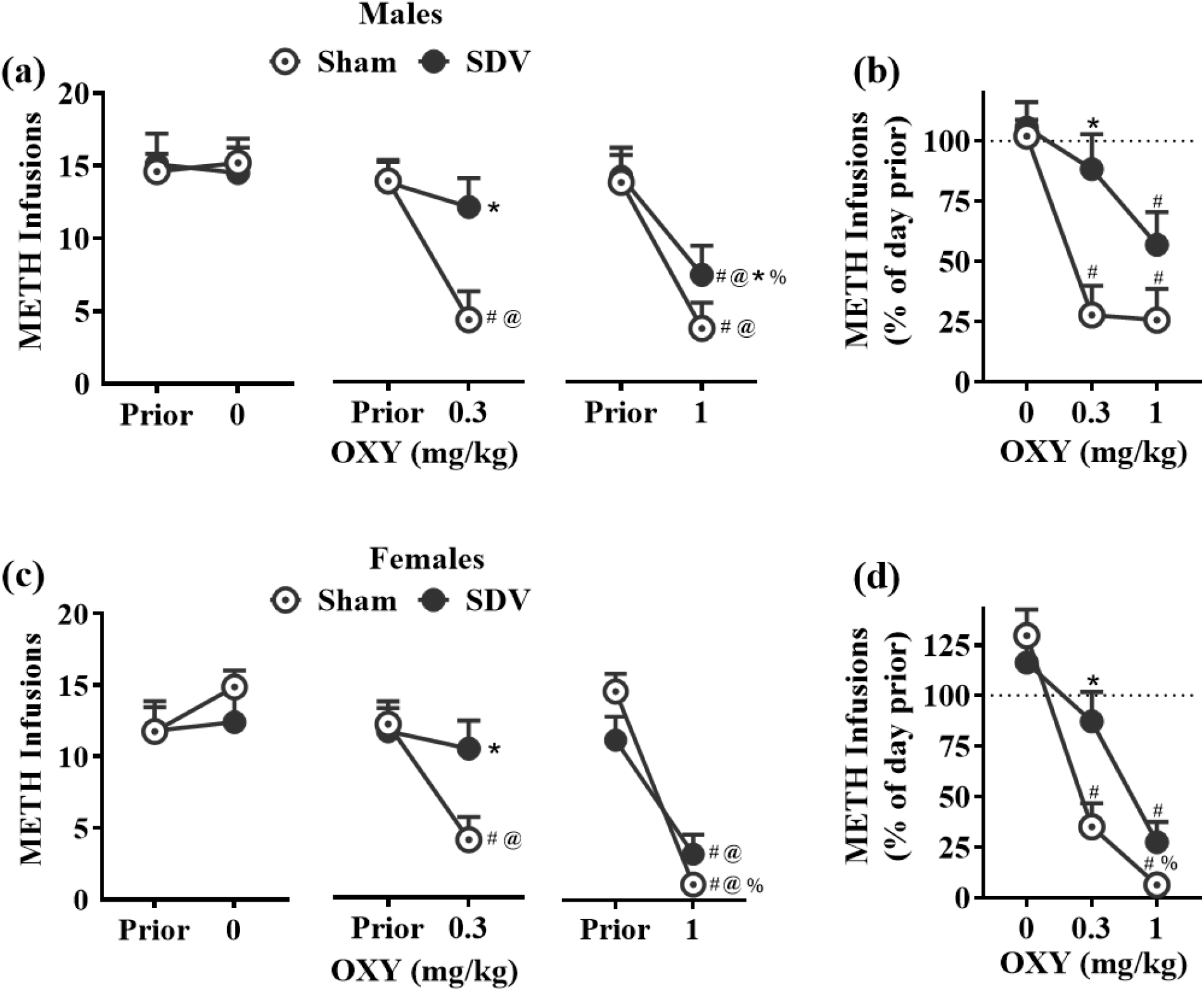
The effects of IP oxytocin treatment on self-administered METH between Sham and SDV rats. Self-administered METH infusions in Sham or SDV rats following 0, 0.3 or 1.0 mg/kg OXY, compared to the day prior to test for (a) males and (c) females. METH infusions normalized to percentage of day prior to the respective test for males (b) and females (d). Data are mean ± SEM. © *p* < 0.05 versus respective day prior. # *p* < 0.05 versus respective 0 mg/kg test. * *p* < 0.05 versus respective Sham control group data point. % *p* < 0.05 between respective 0.3 and 1.0 mg/kg OXY doses.

### Vagotomy attenuated oxytocin inhibition of methamphetamine self-administration in both sexes

Self-administered METH on test days was normalized to intake on the prior day (percent reduction; Figure 2). There was no interaction of sex×surgery×dose on METH intake (F(1,34)=0.295, *p*=0.591), however, there was a significant surgery×dose interaction (F(2,68)=8.267, *p*=0.001). As three levels of dose were included, simple 2×2 interactions were analysed between surgery levels, and within pairs of levels of dose. This revealed significant surgery×dose interactions when comparing 0 vs 0.3 mg/kg (F(1,34)=11.695, *p*=0.002) and 0 vs 1 mg/kg (F(1,34)=5.372, *p*=0.027), whereby oxytocin treatment suppressed METH intake less in SDV rats than in sham operated rats. Interestingly, there was a significant interaction of surgery×dose when comparing 0.3 vs 1 mg/kg (F(1,34)=5.035, *p*=0.031), as SDV attenuated the inhibitory effect of oxytocin on METH intake more so at 0.3 than at 1 mg/kg. There was also a significant interaction of sex×dose on infusions when comparing 0 vs 1 mg/kg (F(1,34)=9.738, *p*=0.004), but not 0 vs 0.3 (F(1,34)=1.304, *p*=0.261), or 0.3 vs 1 mg/kg (F(1,34)=3.456, *p*=0.072), indicating that 1 mg/kg oxytocin was more effective at reducing METH intake in females. Simple contrasts comparing METH infusions between individual levels of dose, sex, and surgery are depicted in Figure 2, and in Supplemental Tables 1-5.

### Extinction of methamphetamine self-administration

Rats decreased their active nose-pokes and locomotor activity over the 14 days of extinction (for all groups, Day 1 vs 14: *p*<0.05), and all rats met standard criteria of <10 active nose-pokes before proceeding to reinstatement tests (Supplemental Figure 1).

### Vagotomy attenuated oxytocin inhibition of cue-induced reinstatement in males but not females

Assessing active nose pokes during cue-induced reinstatement, there were no interactions of sex×surgery×dose (F1,33)=0.975, *p*=0.331; Figure 3) or sex×surgery (F(1,33)=1.115, *p*=0.299), and no main effects of surgery (F(1, 33)=0.062, *p*=0.805) or sex (F(1, 33)=1.297, *p*=0.263). There was a significant main effect of oxytocin dose on active nose-pokes (F(1,33)=39.630, *p*<0.001), and significant interactions of dose×sex (F(1,33)=7.480, *p*=0.010) and dose×surgery (F(1,33)=5.711, *p*=0.023), whereby the inhibitory effects of oxytocin on cue-induced reinstatement were more potent in females, and were attenuated by SDV. To best assess the effects of SDV and oxytocin within each sex, active nose-pokes following 0.3 mg/kg oxytocin were normalised to the vehicle session. This showed that oxytocin suppressed active nose-pokes significantly less in SDV than sham males (*t*(17)=2.208, *p*=0.041), while SDV and sham females were not significantly different (*t*(16)=0.887, *p*=0.388; Fig 5d). Simple contrasts comparing active nose-pokes and locomotor activity between individual levels of dose, sex, and surgery are depicted in figure 3, and in Supplemental Tables 6-9.

**Figure 3.**
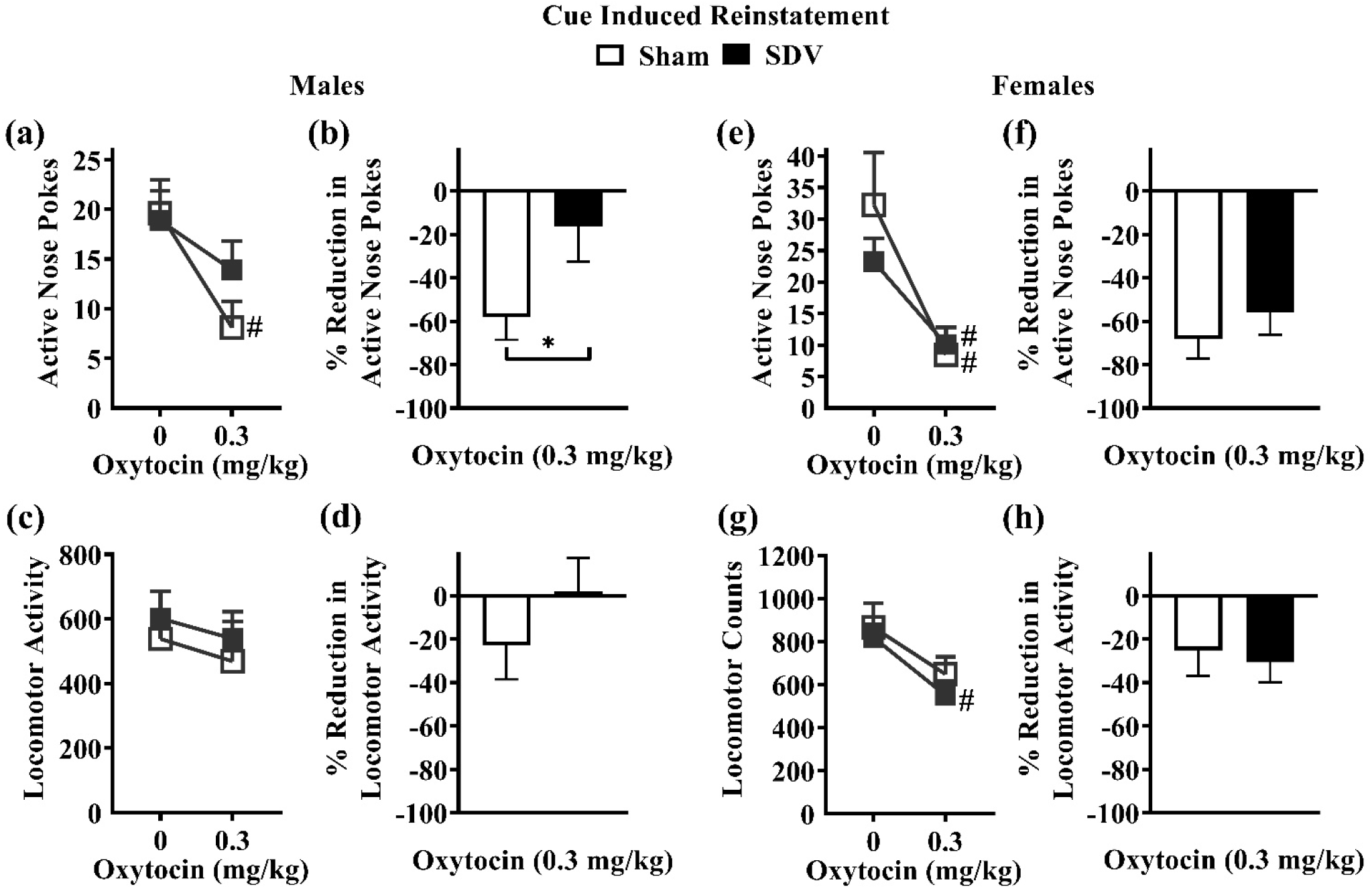
The effects of IP oxytociu treatment on cue-induced reinstatement to METH-seeking behaviours Sham and SDV rats. Cue-induced reinstatement of active nose pokes(a and e), active nose pokes normalised to 0 mg/kg test (b and f), locomotor activity (c and g), locomotor activity normalised to 0 mg/kg test and (d and h), for males (a-d) and females (e-h) respectively. Data are mean ± SEM. # *p* < 0.05 versus respective 0 mg/kg test. * *p* < 0.05 versus respective Sham control group data point.

### Vagotomy attenuated oxytocin inhibition of methamphetamine-primed reinstatement in males but not females

There were significant main effects of sex (F(1,33)=4.658, *p*=0.038) and dose (F(2,33)=20.235, *p*<0.001), but not SDV (F(1,33)=0.011, *p*=0.917) on active nose-pokes made during METH-primed reinstatement (Figure 4). There was a significant surgery×dose interaction (F(2,66)=3.309, *p*=0.043), and as three levels of dose were included, 2×2 interactions were conducted between surgery levels, and within pairs of levels of dose for males and females separately. For males, this revealed a significant surgery×dose interaction when comparing 0 vs 1 mg/kg (F(1,17)=8.529, *p*=0.010), but not 0 vs 0.3 mg/kg (F(1,17)=2.844, *p*=0.110) or 0.3 vs 1 mg/kg (F(1,17)=0.379, *p*=0.546) indicating that 1 mg/kg oxytocin reduced METH-primed reinstatement more so in sham than SDV males. For females, there were no significant surgery×dose interactions when comparing 0 vs 0.3 mg/kg (F(1,16)=0.531, *p*=0.477), 0 vs 1 mg/kg (F(1,16)=0.760, *p*=0.396), or 0.3 vs 1 mg/kg (F(1,16)=2.689, *p*=0.121) indicating that both oxytocin doses reduced METH-primed reinstatement similarly in sham and SDV females. In further support of these interactions, when the effects of oxytocin on active nose-pokes were normalised to the vehicle session, SDV males nose-poked significantly more than shams at both 0.3 mg/kg (*p*=0.039) and 1.0 mg/kg doses (*p*=0.047), while SDV females were not significantly different to sham females at either dose (*p*=0.264 and 0.315 respectively). Simple contrasts comparing active nose-pokes and locomotor activity between individual levels of dose, sex, and surgery are depicted in figure 4, and in Supplemental Tables 10-13.

**Figure 4.**
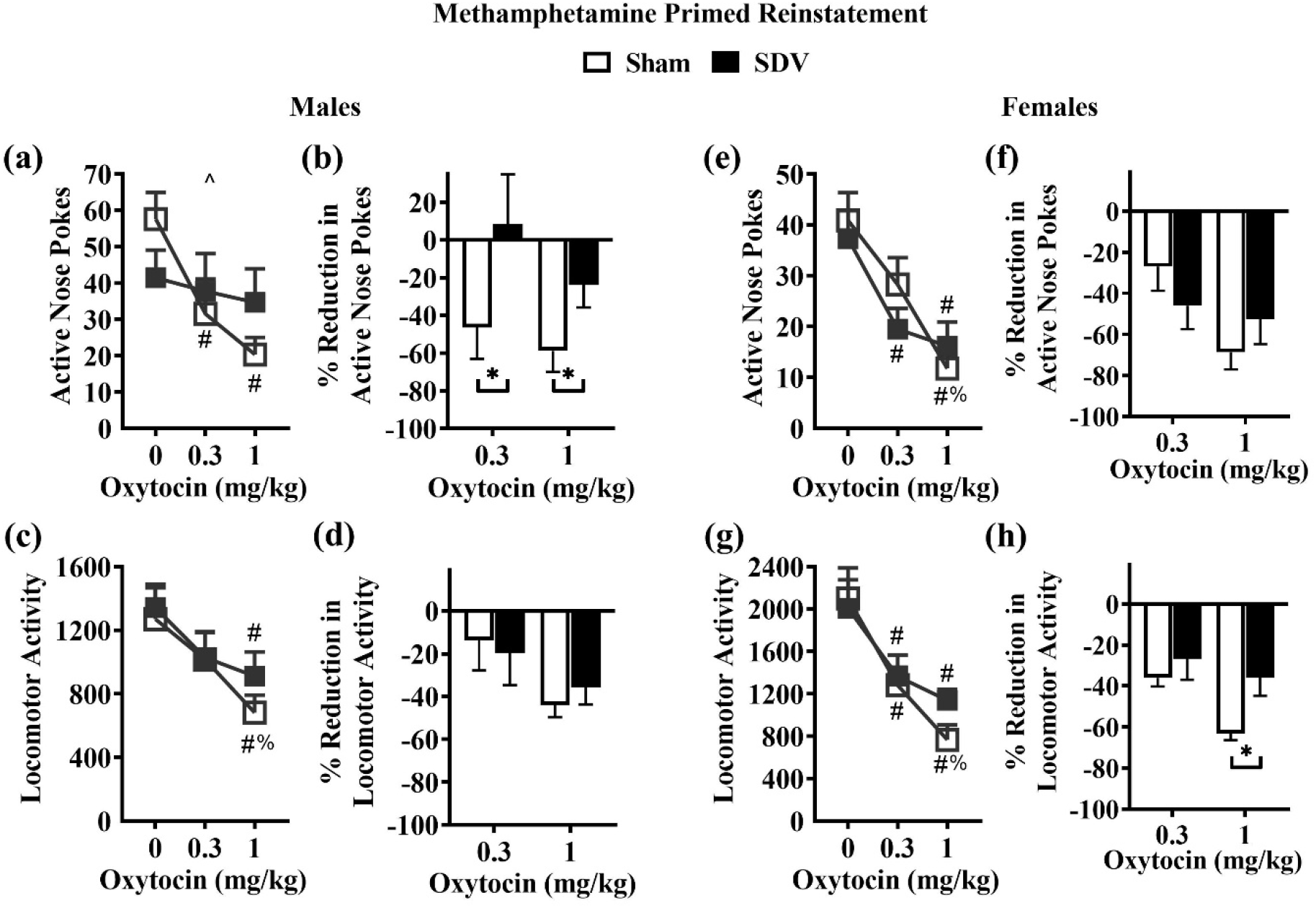
The effects of IP oxytocin trea tment on METH-primed reinstatement to METH­ seeking behaviours Sham and SDV r ats. METH-primed reinstatement of active nose pokes (a and e), active nose pokes normalised to 0 mg/kg test (band f), locomotoractivity (c and g), and locomotor activity normalised to 0 mg/kg test (d and h), for males (a-d) and females (e-h) respectively. Data are mean± SEM. % *p* < 0.05 benveenrespective 0.3 and 1mg/kg OXY doses. # *p* < 0.05 versus respective 0 mg/kg test. * *p* < 0.05 versus respective Sham control group data point. ^ *p* < 0.05 interaction of dose (0 vs 1mg/kg) and surgery (Sham vs SDV).

**Figure 5.**
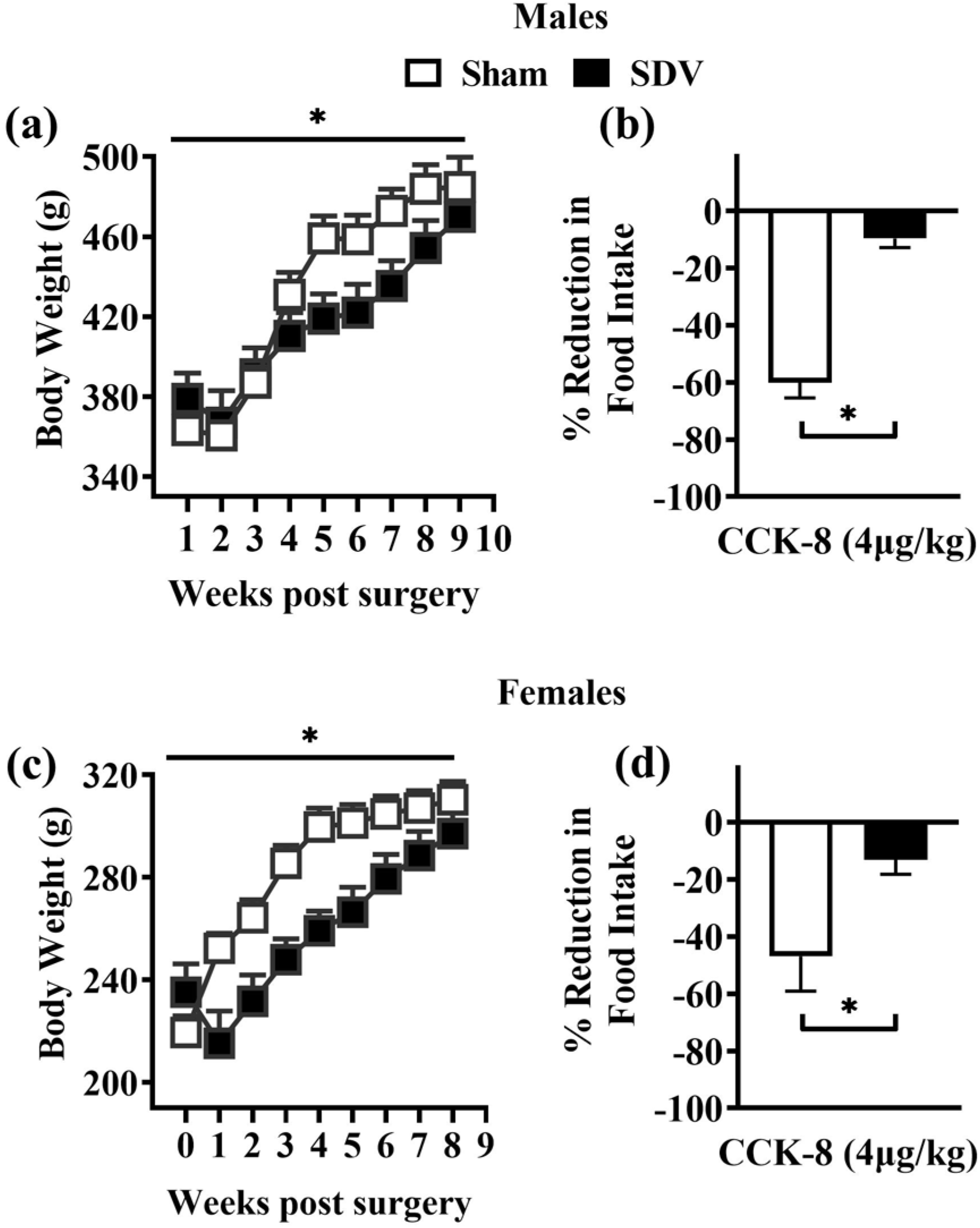
Body weight, and functional validation of successful subdia phragmatic vagotom y by CCK-8 inhibition of feeding in Sham and SDV rats. Body weight over 10 weeks for Sham and SDV males (a) and females (c). Percent reduction of intake of liquid rodent diet following IP injection of CCK-8(4 ug/kg) in Sham and SDV males (b) and females (d). *p* <0.05 between Sham and SDV rats. Data are mean ± SEM

### Vagotomy was functionally confirmed by cholecystokinine-8 suppression of food consumption

There was a significant main effect of surgery (F(1,33)=6.037, *p*=0.019; Figure 7), whereby SDV rats had lower body weight than sham rats, and a significant interaction of surgery×week (F(8,272)=6.524, *p*<0.001), indicating the weight differences dissipated over time. There was no sex×surgery interaction for weight (F(1,33)=0.099, *p*=0.755). CCK-8 treatment reduced feeding by at least 35% in all sham rats, whereas no SDV rats had a reduction of intake greater than 24%. There was a significant effect of surgery (F(1,33)=37.902, *p*<0.001), whereby feeding was reduced in sham but not SDV rats. There was no interaction of sex×surgery (F(1,33)=1.486, *p*=0.234), and no main effect of sex (F(1,33)=0.516, *p*=0.478), on reduction of food intake following CCK-8 treatment.

## Discussion

Here we present the novel finding that the inhibitory effect of peripherally administered oxytocin on METH addiction-like behaviours depends on subdiaphragmatic vagal signaling. Specifically, the effect of a low dose of oxytocin to inhibit METH self-administration were absent in male and female rats which had undergone subdiaphragmatic vagotomy, yet persisted in sham controls. Conversely, the inhibitory effect of a higher dose of oxytocin on METH intake were only partially attenuated by vagotomy in males, and were vagal-independent in females. After a period of extinction, vagotomy reduced the inhibitory effect of oxytocin treatment on cue-induced and METH-primed reinstatement to METH-seeking behaviours in males, but not females. These data provide clear evidence for a role of vagal signaling in the anti-addiction effect of oxytocin treatment and suggest that sex-dependent oxytocin-vagal mechanisms emerge following METH withdrawal.

Oxytocin administered at 0.3 mg/kg potently reduced METH self-administration, which was almost entirely prevented by SDV in both sexes. However, at the 1 mg/kg oxytocin dose, this vagal dependency was less pronounced. Sham females were also more sensitive to this higher dose compared to sham males. One explanation for this dose-dependent involvement of the vagus nerve for the effect of oxytocin is increased penetration of the blood-brain barrier at higher doses of oxytocin^47^. Recent studies in oxytocin-null mice, which cannot produce oxytocin, demonstrate that peripheral oxytocin injection results in a substantial increase in brain oxytocin levels^48,49^. As such, it is possible that the brain concentrations of oxytocin achieved by the low dose are insufficient to inhibit METH self-administration in SDV rats. Rather, suppression of METH intake by this low dose in shams may be achieved through stimulation of an ascending vagal pathway to the brain. In contrast, the high oxytocin dose may result in brain concentrations of oxytocin which *are* sufficient to inhibit METH seeking, despite the lack of vagal signaling. Both sexes were similar with regard to the vagal-dependent effects of IP oxytocin on suppressing METH-intake. These data clearly implicate the vagus nerve as a crucial target for IP oxytocin in reducing METH intake, and yet are not inconsistent with observations of behaviorally relevant blood-brain passage of peripherally administered oxytocin, particularly at higher doses.

Following extinction of drug-seeking and withdrawal from METH, the role of the vagus nerve in mediating the effects of oxytocin on METH behaviours appears to be more complex. In males, 0.3 mg/kg oxytocin reduced cue-induced reinstatement, which was partially attenuated by vagotomy, and both the effect of oxytocin and the interference by SDV seemed independent of a locomotor effect. For METH-primed reinstatement, 0.3 and 1.0 mg/kg oxytocin potently reduced METH-primed reinstatement in shams but had no effect in SDV males, which was also separable from the effect of oxytocin on METH-induced locomotor activity. These results indicate that despite undergoing extinction and withdrawal, the vagus nerve mediates the inhibitory effects of peripheral oxytocin treatment on METH-seeking behaviours in males.

In females, 0.3 mg/kg oxytocin potently reduced cue-induced reinstatement, and 0.3 and 1.0 mg/kg oxytocin reduced METH-primed reinstatement, in both sham and SDV females. These effects of oxytocin on reinstatement, and interference by SDV are largely independent of an effect on locomotor activity, although the 1 mg/kg dose attenuated METH-induced locomotor activity less in SDV than in sham females. Surprisingly, the female reinstatement data is in stark contrast to the almost entire vagal dependency of the effects of 0.3 mg/kg oxytocin on METH self-administration, and slight mediation of the 1.0 mg/kg dose by SDV in these same rats. It is also in contrast to the enduring vagal dependency of oxytocin’s effects seen in male SDV rats.

The divergent interactions of SDV, oxytocin, and sex between self-administration and reinstatement are challenging to interpret, due to the lack of female subjects included in oxytocin, METH, and vagal research. However, there are several possible explanations for the present sex differences. Firstly, there is evidence that expression of oxytocin receptors is altered by chronic METH self-administration and extinction in male rodents^50,51^. As females and males have non-identical patterns of oxytocin receptor expression^52^ it is possible that METH- and extinction-induced plasticity of oxytocin receptors may result in an augmented sensitivity to the concentration of IP oxytocin which crosses the BBB in SDV females. Alternatively, METH self-administration causes BBB injury^53^, which may result in greater penetration of IP oxytocin into the brain. Indeed, the mechanism by which oxytocin is transported into the brain has recently been identified^48^, and this system may be affected by METH exposure^54^. However, the effect of METH exposure and withdrawal on oxytocin receptors or BBB integrity has mostly been studied in male subjects, so it is unclear whether there are sex-dependent effects of METH on either of these systems. Lastly, the effects of VNS in addiction models have also only been explored in males^41,42^, rendering it unknown whether VNS can achieve similar anti-addiction effects in females.

It is unlikely that the altered vagal dependency of oxytocin treatment observed in females following extinction could be due to regeneration of vagal nerve endings over the 3 weeks separating self-administration and reinstatement tests. Vagal regeneration can occur after ~5 months post-SDV in rats, however even after this time it is estimated that only ~7-39% of fibres regenerate^55^. Given the 3-week timespan separating our vagal-dependent and vagal-independent effects of oxytocin in females, regeneration is unlikely to explain our data. Indeed, if this effect was due to vagal regeneration, then the enduring vagal dependency of oxytocin in males would suggest profound sex differences in vagal regrowth. We are not aware of any literature to support sex-dependent regeneration of peripheral nerves to this extent. Regardless, functional vagotomy was confirmed in both sexes after reinstatement tests by administering the satiety-signaling peptide CCK-8, which inhibits feeding through subdiaphragmatic vagal signaling^56^. This test at the very least confirms that vagal afferents sensitive to CCK-8 were non-functional. As oxytocin has been shown to stimulate a similar population of nodose ganglion neurons as CCK-8^32^, we can be confident that the sex-dependent mediation of oxytocin treatment on reinstatement by SDV is not due to vagal regeneration.

Activation of the vagus nerve by peripheral oxytocin treatment produces a cascade of effects along a central projecting pathway. The first step of this pathway involves vagal innervation of the NTS. Peripheral oxytocin injection increases neuronal activity within the NTS, which is entirely prevented by SDV^32^. Noradrenergic neurons in the NTS project, via polysynaptic pathways involving the locus coeruleus^57^, to widespread structures with relevance to METH addiction, including the extended amygdala complex which mediates activity of the nucleus accumbens^58^, a crucial addiction substrate. It is possible that IP oxytocin modulates activity at these addiction-relevant structures via a vagal-NTS pathway.

Stimulation of this vagal-NTS pathway may also produce anti-addiction effects through activation of the endogenous oxytocin system. Noradrenergic neurons in the NTS also project to the PVN^59^ and can activate oxytocin cells^60^. Indeed, lesions to the NTS markedly reduce reactivity of PVN oxytocin cells in response to a peripheral inflammatory challenge^29^. Stimulation of oxytocin neurons via a vagal-NTS-PVN pathway is further supported by recent demonstration that SDV prevents IP oxytocin from increasing activity of NTS-PVN projecting neurons, and prevents oxytocin release in the PVN^61^. Oxytocin-secreting cells in the PVN project to a variety of structures implicated in addiction. For example, stimulation of oxytocin fibres in the central amygdala causes local GABAergic inhibition^62^, an effect which has been shown to reduce cue-induced reinstatement to METH-seeking^63^. Therefore, IP oxytocin may stimulate endogenous release of oxytocin at sites which govern drug seeking, via this vagal-NTS-PVN pathway.

The vagal-independent effects of the high oxytocin dose may be explained by circulating oxytocin stimulating other peripheral structures. For example, high doses of oxytocin may avoid the vagal-dependent access to this NTS-PVN pathway through stimulation of the BBB-deficient area postrema, which mediates the ability of the NTS to stimulate PVN oxytocin release^64^. Alternatively, high levels of peripherally circulating oxytocin could reach the nodose ganglion, activating the cell bodies of the vagi which project to the NTS. Such a vagal- or nodose-dependent effect has been shown in the transmission of peripheral immune signals to the brain^65^, and may explain our findings.

These findings have implications for the design and interpretation of clinical studies, and for the applicability of oxytocin as a pharmacotherapy for METH-use disorders. Firstly, it is crucial that sex is included as a biological variable. Currently, the only registered trial investigating the efficacy of oxytocin on METH addiction outcomes is in treatment seeking HIV-positive males^22^. As such, it should not be presumed that the results of this trial will inform the suitability of oxytocin as a therapy for females, who according to our findings, and others^43^, may be more sensitive to oxytocin treatment, and may have non-identical peripheral-to-central signaling pathways for oxytocin. Furthermore, as the vagal-dependent effects of peripherally administered oxytocin may change throughout withdrawal, oxytocin should be explored at timepoints encompassing current use, and into abstinence. Lastly, as intranasal delivery of oxytocin in the clinic gains both momentum and controversy^23^, the differences between intranasal and IP administration must be considered. Unlike IP, IN oxytocin does not increase activity in the mouse NTS^66^, which may preclude the therapeutic benefits of stimulating this vagal-mediated pathway. Furthermore, IP oxytocin produces substantially higher plasma levels of oxytocin than IN^67^, which may contribute to the vagal-dependent effects of IP administration. However, as IN oxytocin does elevate plasma levels of oxytocin, it would be intriguing to investigate whether the behavioural effects of IN oxytocin are prevented by SDV. The translation of oxytocin from IP in rodents, to IN in humans may depend on how important this oxytocin-stimulated vagal-brain pathway is for treating addictions in humans, and whether IN oxytocin can stimulate this pathway.

In conclusion, we discovered that the well documented preclinical effects of peripherally administered oxytocin in reducing METH self-administration and reinstatement to METH-seeking behaviours is largely dependent upon the subdiaphragmatic vagus nerve. Sex differences in this vagal dependency of oxytocin’s effects emerged after withdrawal, which has implications for the clinical utility of oxytocin. Lastly, the growing momentum towards intranasal delivery of oxytocin to bypass the blood-brain barrier should be considered in light of these findings, which indicate that in rats, a considerable amount of the therapeutic effects of oxytocin for treating METH addiction is mediated by the vagus nerve. As such, drugs or implants which target oxytocin-sensitive vagal pathways may prove to be valuable therapies for METH addiction.

## Acknowledgements

This research was supported by the Department of Psychology, Macquarie University. NAE and PAC were recipients of Australia Postgraduate Award scholarships. NAE, AJT, and JLC performed surgeries. NAE and SJB carried out post-operative care. NAE carried out self-administration experiments. NAE and PAC carried out feeding experiments. NAE conceived the study. NAE, SJB, and JLC designed the experiments, and AJT designed the surgical procedure. NAE conducted data analysis and wrote the manuscript. All authors critically reviewed the content and approved the final version before submission. We thank Justin Clarke and Ronny Eidels for their assistance with designing the surgical procedures, Christine Sutter for her all-encompassing support, and Virginia Williams, Carlie Crawford, Julie Reynolds, and the animal technicians for their daily support.

## References

1. Darke, S., Kaye, S., McKETIN, R. & Duflou, J. Major physical and psychological harms of methamphetamine use. Drug Alcohol Rev. 27, 253–262 (2008).

2. A Meta-Analytic Review of Psychosocial Interventions for Substance Use Disorders | American Journal of Psychiatry. https://ajp.psychiatryonline.org/doi/full/10.1176/appi.ajp.2007.06111851#.

3. Forray, A. & Sofuoglu, M. Future pharmacological treatments for substance use disorders. Br. J. Clin. Pharmacol. 77, 382–400 (2014).

4. Morley, K. C., Cornish, J. L., Faingold, A., Wood, K. & Haber, P. S. Pharmacotherapeutic agents in the treatment of methamphetamine dependence. Expert Opin. Investig. Drugs 26, 563–578 (2017).

5. Ballester, J., Valentine, G. & Sofuoglu, M. Pharmacological treatments for methamphetamine addiction: current status and future directions. Expert Rev. Clin. Pharmacol. 10, 305–314 (2017).

6. McGregor, I. S. & Bowen, M. T. Breaking the loop: Oxytocin as a potential treatment for drug addiction. Horm. Behav. 61, 331–339 (2012).

7. Sarnyai, Z. & Kovács, G. L. Oxytocin in learning and addiction: From early discoveries to the present. Pharmacol. Biochem. Behav. 119, 3–9 (2014).

8. Baracz, S. J. & Cornish, J. L. The neurocircuitry involved in oxytocin modulation of methamphetamine addiction. Front. Neuroendocrinol. 43, 1–18 (2016).

9. Lee, M. R. & Weerts, E. M. Oxytocin for the treatment of drug and alcohol use disorders. Behav. Pharmacol. 27, 640–648 (2016).

10. Bowen, M. T. & Neumann, I. D. Rebalancing the Addicted Brain: Oxytocin Interference with the Neural Substrates of Addiction. Trends Neurosci. 40, 691–708 (2017).

11. Carson, D. S., Cornish, J. L., Guastella, A. J., Hunt, G. E. & McGregor, I. S. Oxytocin decreases methamphetamine self-administration, methamphetamine hyperactivity, and relapse to methamphetamine-seeking behaviour in rats. Neuropharmacology 58, 38–43 (2010).

12. Cox, B. M. et al. Oxytocin Acts in Nucleus Accumbens to Attenuate Methamphetamine Seeking and Demand. Biol. Psychiatry 81, 949–958 (2017).

13. Everett, N., Baracz, S. & Cornish, J. Oxytocin treatment in the prelimbic cortex reduces relapse to methamphetamine-seeking and is associated with reduced activity in the rostral nucleus accumbens core. Pharmacol. Biochem. Behav. 183, 64–71 (2019).

14. Bernheim, A., Leong, K.-C., Berini, C. & Reichel, C. M. Antagonism of mGlu2/3 receptors in the nucleus accumbens prevents oxytocin from reducing cued methamphetamine seeking in male and female rats. Pharmacol. Biochem. Behav. 161, 13–21 (2017).

15. Everett, N. A., McGregor, I. S., Baracz, S. J. & Cornish, J. L. The role of the vasopressin V1A receptor in oxytocin modulation of methamphetamine primed reinstatement. Neuropharmacology 133, 1–11 (2018).

16. Baracz, S. J., Everett, N. A., McGregor, I. S. & Cornish, J. L. Oxytocin in the nucleus accumbens core reduces reinstatement of methamphetamine-seeking behaviour in rats. Addict. Biol. 21, 316–325 (2016).

17. Baracz, S. J., Everett, N. A. & Cornish, J. L. The Involvement of Oxytocin in the Subthalamic Nucleus on Relapse to Methamphetamine-Seeking Behaviour. PLOS ONE 10, e0136132 (2015).

18. The effect of chronic oxytocin treatment during abstinence from methamphetamine self-administration on incubation of craving, reinstatement, and anxiety | Neuropsychopharmacology. https://www.nature.com/articles/s41386-019-0566-6.

19. McRae-Clark, A. L., Baker, N. L., Maria, M. M.-S. & Brady, K. T. Effect of oxytocin on craving and stress response in marijuana-dependent individuals: a pilot study. Psychopharmacology (Berl.) 228, 623–631 (2013).

20. Pedersen, C. A. et al. Intranasal Oxytocin Blocks Alcohol Withdrawal in Human Subjects. Alcohol. Clin. Exp. Res. 37, 484–489 (2013).

21. Hansson, A. C. et al. Oxytocin Reduces Alcohol Cue-Reactivity in Alcohol-Dependent Rats and Humans. Neuropsychopharmacology 43, 1235–1246 (2018).

22. Stauffer, C. S. et al. Oxytocin-enhanced motivational interviewing group therapy for methamphetamine use disorder in men who have sex with men: study protocol for a randomized controlled trial. Trials 20, 145 (2019).

23. Leng, G. & Ludwig, M. Intranasal Oxytocin: Myths and Delusions. Biol. Psychiatry 79, 243–250 (2016).

24. Mens, W. B. J., Witter, A. & Van Wimersma Greidanus, T. B. Penetration of neurohypophyseal hormones from plasma into cerebrospinal fluid (CSF): Half-times of disappearance of these neuropeptides from CSF. Brain Res. 262, 143–149 (1983).

25. Born, J. et al. Sniffing neuropeptides: a transnasal approach to the human brain. Nat. Neurosci. 5, 514–516 (2002).

26. Kimura, T. et al. Molecular regulation of the oxytocin receptor in peripheral organs. J. Mol. Endocrinol. 109–115 (2003) doi:10.1677/jme.0.0300109.

27. Prechtl, J. C. & Powley, T. L. The fiber composition of the abdominal vagus of the rat. Anat. Embryol. (Berl.) 181, 101–115 (1990).

28. Delfs, J. M., Zhu, Y., Druhan, J. P. & Aston-Jones, G. S. Origin of noradrenergic afferents to the shell subregion of the nucleus accumbens: anterograde and retrograde tract-tracing studies in the rat. Brain Res. 806, 127–140 (1998).

29. Buller, K., Xu, Y., Dayas, C. & Day, T. Dorsal and Ventral Medullary Catecholamine Cell Groups Contribute Differentially to Systemic Interleukin-1β-Induced Hypothalamic Pituitary Adrenal Axis Responses. Neuroendocrinology 73, 129–138 (2001).

30. Stock, S. & Uvnäs-Moberg, K. Increased plasma levels of oxytocin in response to afferent electrical stimulation of the sciatic and vagal nerves and in response to touch and pinch in anaesthetized rats. Acta Physiol. Scand. 132, 29–34 (1988).

31. Cunningham, J. T., Mifflin, S. W., Gould, G. G. & Frazer, A. Induction of c-Fos and ∆FosB Immunoreactivity in Rat Brain by Vagal Nerve Stimulation. Neuropsychopharmacology 33, 1884–1895 (2008).

32. Iwasaki, Y. et al. Peripheral oxytocin activates vagal afferent neurons to suppress feeding in normal and leptin-resistant mice: a route for ameliorating hyperphagia and obesity. Am. J. Physiol.-Regul. Integr. Comp. Physiol. 308, R360–R369 (2014).

33. Gao, X. et al. Presence of functional vasopressin V1 receptors in rat vagal afferent neurones. Neurosci. Lett. 145, 79–82 (1992).

34. Yuan, H. & Silberstein, S. D. Vagus Nerve and Vagus Nerve Stimulation, a Comprehensive Review: Part II. Headache J. Head Face Pain 56, 259–266 (2016).

35. De Cagna, F. et al. The Role of Intranasal Oxytocin in Anxiety and Depressive Disorders: A Systematic Review of Randomized Controlled Trials. Clin. Psychopharmacol. Neurosci. 17, 1–11 (2019).

36. Rush, A. J. et al. Vagus Nerve Stimulation for Treatment-Resistant Depression: A Randomized, Controlled Acute Phase Trial. Biol. Psychiatry 58, 347–354 (2005).

37. Neumann, I. D. & Slattery, D. A. Oxytocin in General Anxiety and Social Fear: A Translational Approach. Biol. Psychiatry 79, 213–221 (2016).

38. George, M. S. et al. A pilot study of vagus nerve stimulation (VNS) for treatment-resistant anxiety disorders. Brain Stimulat. 1, 112–121 (2008).

39. Olff, M., Langeland, W., Witteveen, A. & Denys, D. A Psychobiological Rationale for Oxytocin in the Treatment of Posttraumatic Stress Disorder. CNS Spectr. 15, 522–530 (2010).

40. Noble, L. J. et al. Effects of vagus nerve stimulation on extinction of conditioned fear and post-traumatic stress disorder symptoms in rats. Transl. Psychiatry 7, e1217–e1217 (2017).

41. Liu, H. et al. Vagus nerve stimulation inhibits heroin-seeking behavior induced by heroin priming or heroin-associated cues in rats. Neurosci. Lett. 494, 70–74 (2011).

42. Childs, J. E., DeLeon, J., Nickel, E. & Kroener, S. Vagus nerve stimulation reduces cocaine seeking and alters plasticity in the extinction network. Learn. Mem. 24, 35–42 (2017).

43. Cox, B. M., Young, A. B., See, R. E. & Reichel, C. M. Sex differences in methamphetamine seeking in rats: Impact of oxytocin. Psychoneuroendocrinology 38, 2343–2353 (2013).

44. Dluzen, D. E. & Liu, B. Gender differences in methamphetamine use and responses: A review. Gend. Med. 5, 24–35 (2008).

45. Mravec, B., Ondicova, K., Tillinger, A. & Pecenak, J. Subdiaphragmatic vagotomy enhances stress-induced epinephrine release in rats. Auton. Neurosci. 190, 20–25 (2015).

46. Zhang, J. & Ritter, R. C. Circulating GLP-1 and CCK-8 reduce food intake by capsaicin-insensitive, nonvagal mechanisms. Am. J. Physiol.-Regul. Integr. Comp. Physiol. 302, R264–R273 (2012).

47. Bowen, M. T. Does peripherally administered oxytocin enter the brain? Compelling new evidence in a long-running debate. Pharmacol. Res. 146, 104325 (2019).

48. Yamamoto, Y. et al. Vascular RAGE transports oxytocin into the brain to elicit its maternal bonding behaviour in mice. Commun. Biol. 2, 1–13 (2019).

49. Smith, A. S., Korgan, A. C. & Young, W. S. Oxytocin delivered nasally or intraperitoneally reaches the brain and plasma of normal and oxytocin knockout mice. Pharmacol. Res. 146, 104324 (2019).

50. Baracz, S. J. et al. Chronic Methamphetamine Self-Administration Dysregulates Oxytocin Plasma Levels and Oxytocin Receptor Fibre Density in the Nucleus Accumbens Core and Subthalamic Nucleus of the Rat. J. Neuroendocrinol. 28, (2016).

51. Georgiou, P. et al. Methamphetamine abstinence induces changes in μ-opioid receptor, oxytocin and CRF systems: Association with an anxiogenic phenotype. Neuropharmacology 105, 520–532 (2016).

52. Smith, C. J. W. et al. Age and sex differences in oxytocin and vasopressin V1a receptor binding densities in the rat brain: focus on the social decision-making network. Brain Struct. Funct. 222, 981–1006 (2017).

53. Ramirez, S. H. et al. Methamphetamine Disrupts Blood–Brain Barrier Function by Induction of Oxidative Stress in Brain Endothelial Cells. J. Cereb. Blood Flow Metab. 29, 1933–1945 (2009).

54. Treweek, J. B., Dickerson, T. J. & Janda, K. D. Drugs of abuse that mediate advanced glycation end product formation: A chemical link to disease pathology. Acc. Chem. Res. 42, 659–669 (2009).

55. Phillips, R. J., Baronowsky, E. A. & Powley, T. L. Regenerating vagal afferents reinnervate gastrointestinal tract smooth muscle of the rat. 22.

56. Abdominal vagotomy blocks the satiety effect of cholecystokinin in the rat | Science. https://science.sciencemag.org/content/213/4511/1036?casa_token=PQ8D1MeDzfEAAAAA:H6LgcaUUXRU3RgqnxVVC5ngzqDTPLsqGJDPIiHojRuwDLndXL5_RVejhFuSo4UsF4AiQyRg-x96aKho.

57. Hassert, D. L., Miyashita, T. & Williams, C. L. The effects of peripheral vagal nerve stimulation at a memory-modulating intensity on norepinephrine output in the basolateral amygdala. Behav. Neurosci. 118, 79–88 (2004).

58. Aston-Jones, G. & Kalivas, P. W. BRAIN NOREPINEPHRINE REDISCOVERED IN ADDICTION RESEARCH. Biol. Psychiatry 63, 1005–1006 (2008).

59. Cunningham, E. T. & Sawchenko, P. E. Anatomical specificity of noradrenergic inputs to the paraventricular and supraoptic nuclei of the rat hypothalamus. J. Comp. Neurol. 274, 60–76 (1988).

60. Onaka, T., Luckman, S. M., Antonijevic, I., Palmer, J. R. & Leng, G. Involvement of the noradrenergic afferents from the nucleus tractus solitarii to the supraoptic nucleus in oxytocin release after peripheral cholecystokinin octapeptide in the rat. Neuroscience 66, 403–412 (1995).

61. Iwasaki, Y. et al. Relay of peripheral oxytocin to central oxytocin neurons via vagal afferents for regulating feeding. Biochem. Biophys. Res. Commun. 519, 553–558 (2019).

62. Knobloch, H. S. et al. Evoked Axonal Oxytocin Release in the Central Amygdala Attenuates Fear Response. Neuron 73, 553–566 (2012).

63. Li, X., Zeric, T., Kambhampati, S., Bossert, J. M. & Shaham, Y. The Central Amygdala Nucleus is Critical for Incubation of Methamphetamine Craving. Neuropsychopharmacology 40, 1297–1306 (2015).

64. Carter, D. A. & Lightman, S. L. A role for the area postrema in mediating cholecystokinin-stimulated oxytocin secretion. Brain Res. 435, 327–330 (1987).

65. Hosoi, T., Okuma, Y., Matsuda, T. & Nomura, Y. Novel pathway for LPS-induced afferent vagus nerve activation: Possible role of nodose ganglion. Auton. Neurosci. 120, 104–107 (2005).

66. Maejima, Y. et al. Nasal oxytocin administration reduces food intake without affecting locomotor activity and glycemia with c-Fos induction in limited brain areas. Neuroendocrinology 101, 35–44 (2015).

67. Neumann, I. D., Maloumby, R., Beiderbeck, D. I., Lukas, M. & Landgraf, R. Increased brain and plasma oxytocin after nasal and peripheral administration in rats and mice. Psychoneuroendocrinology 38, 1985–1993 (2013).

