## Supplemental for "The vagus nerve mediates the suppressing effects of peripherally administered oxytocin on methamphetamine self-administration and seeking in rats"

**Supplemental Methods**

*Pre-operative Care*

Prior to subdiaphragmatic vagotomy surgery, it is essential that rats are transitioned to a nutritionally complete *liquid* diet (Teklad Diet 09215). Additionally, as the vagotomy procedure prevents the pyloric sphincter from passing solid matter from the stomach to the intestines, it is vital that the rats do not consume any solid matter. In our experience, bedding made from corn cob or absorbent fibers will rapidly absorb any spilled liquid food, which the rats will then readily eat, leading to stomach distension. To avoid this entirely, we used recycled paper bedding (1 ply, 5-6 layers), changed fresh daily. We encourage all researchers to use this bedding option, alongside a liquid diet, for long-term behavioural experiments following vagotomy.

*Subdiaphragmatic Vagotomy Surgery*

Rats were anaesthetised using isoflurane gas (3% in 2L/min O2), and received a pre-operative injection of the non-steroidal anti-inflammatory analgesic carprofen (5 mg/kg subcutaneous). The fur was then clipped from the centre of the back, posterior to the shoulder blades, under the left side of the neck, above the jugular vein, and along the left side of the abdomen, spanning from the distal rib cage to the hip. Each clipped area was then thoroughly swabbed with iodine solution. Rats were placed in the supine position and a 5 cm incision was made from the bottom of the left rib cage towards the left hip. The abdominal cavity was held open using blunt surgical retractors wrapped in gauze dampened with saline. Gentle traction was placed on the rostral aspect of the stomach, to expose the esophagus. Additional retractors may be needed to ensure liver lobes are not damaged. Then, using a surgical microscope (Zeiss OPMI VISU 200 S8), the bilateral vagi were identified on the ventral and dorsal surfaces of the esophagus. Nerve hooks were used to isolate and gently strip the vagi from the esophagus, taking care not to damage the surrounding tissues. A section of each nerve was removed by cutting at the most anterior point of the nerve where it enters the diaphragm, and at the most posterior aspect where it approaches the stomach. For sham surgeries, the vagi were isolated and manipulated, but not cut. The abdominal wall muscle was then sutured using absorbable vicryl (J311H, Ethicon), and the skin was closed with surgical staples.

*Jugular Vein Catheterisation Surgery*

All animals then underwent surgery for implantation of a chronic indwelling catheter in the jugular vein for intravenous methamphetamine self-administration procedures. Prior to surgery, catheters were constructed from 14 cm of silastic tubing (0.33 mm internal diameter, 0.64 mm outer diameter, Dow Corning Australia Ptd Ltd, North Ryde, NSW, Australia) attached to a right angled 26 gauge cannula (Plastics One, Wallingford, Connecticut, USA), and mounted to a 2x2cm polypropylene mesh circle (Small Parts, Lexington, Kentucky, USA). On the day of surgery, catheters were disinfected by soaking them in chlorhexidine with alcohol for 30 minutes. A small incision was then made on the left side of the neck (approximately 1 cm in length) and a second right of centre on the back (approximately 3 cm). The mesh back mount was then inserted underneath the skin on the back of the rat, with the exit of the right-angled cannula passing through a small hole on the back (2 mm diameter). The catheter (which has been flushed with sterile saline to ensure there was no remaining chlorhexidine with alcohol solution in it) was then passed under the skin from the back through to the neck incision with the help of closed toothed forceps. The left jugular vein was then tied off and the catheter tubing inserted for 3.5cm towards the heart and secured with surgical thread. The neck and back incision was then cleaned using sterile saline solution and closed with surgical sutures. Catheters were flushed with the antibiotic cephazolin sodium (0.2 ml of 100 mg/ml) and the anticoagulant heparin (60 IU in 0.2ml).

*Post-surgery Care*

Following surgery, the abdominal, neck, and back surgical sites were cleaned with betadine ointment, and the rats were recovered in a heating chamber and then single housed for two days. Rats received subcutaneous treatment with the analgesic carprofen prior to surgery, as well as for 2 days after surgery. To maintain catheter patency, rats received intravenous infusions (0.2ml) with the antibiotic cephazolin during surgery, and then every day for the rest of the experiment. After three days of post-operative care, heparin was added to the daily intravenous infusion to maintain catheter patency. Although all animals were on nutritionally complete liquid diet, supplemented with highly palatable sweetened condensed milk, we also provided rats with HydroGel and DietGel 76A (ClearH_2_O, ME, USA) in the home cage for this post-operative period.

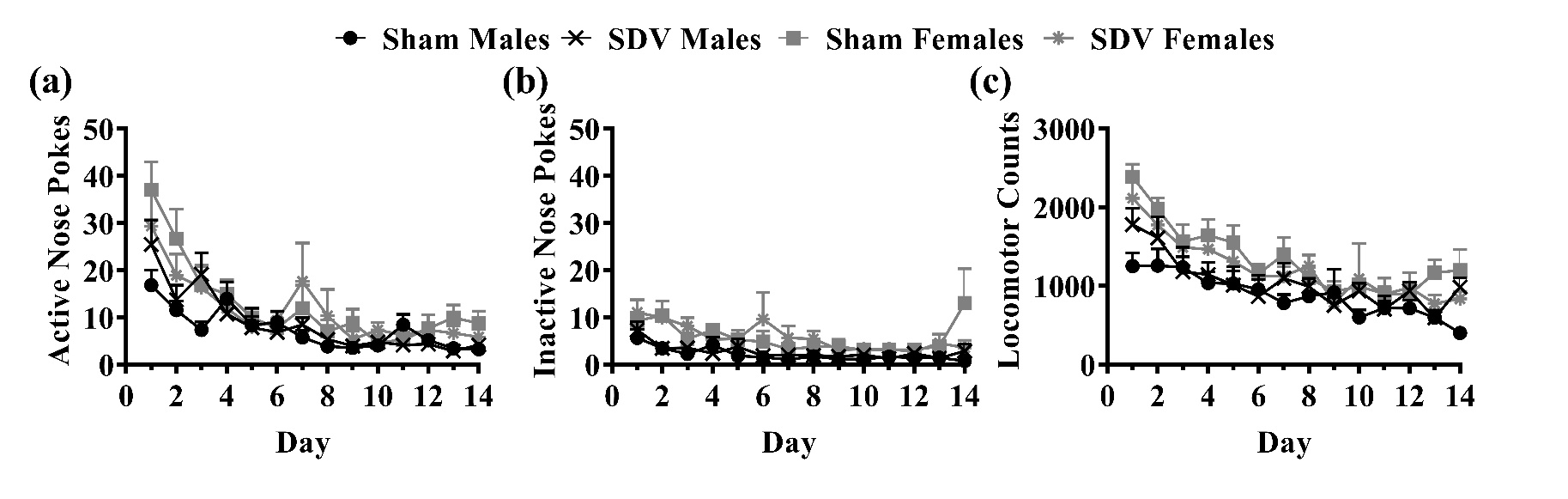
**Supplemental Results**

**Supplemental Figure 1.** Number of (a) active nose pokes, (b) inactive nose pokes, and (c) locomotor activity across 14 days of extinction in sham operated or SDV male and female rats. Data are mean ± SEM.

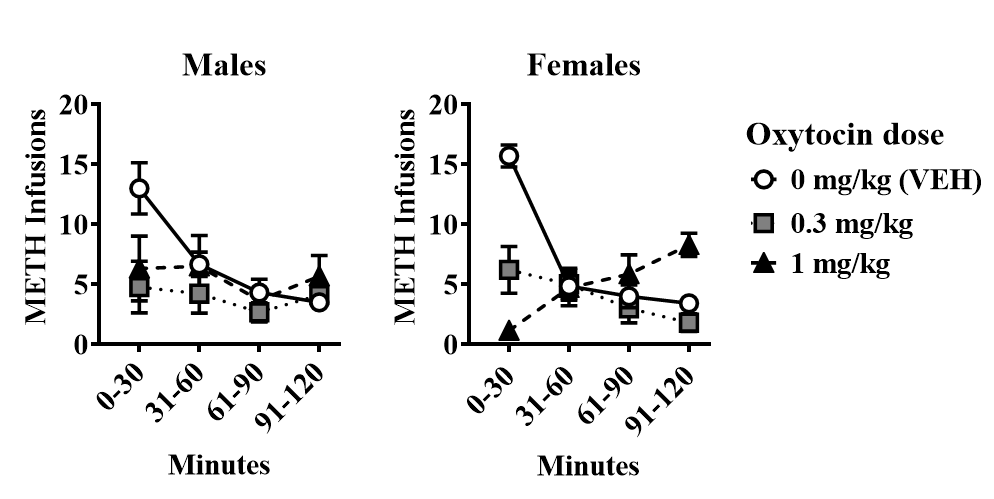

**Supplemental Figure 2.** Oxytocin administered 30-minutes prior to the METH self-administration session causes a time-dependent suppression of METH intake in sham-operated rats (dose×time *p*<0.05). For both sexes, the effects of oxytocin are pronounced in the first 30-minutes of the session but return to vehicle-like levels after that time-point. Therefore, comparisons between sham and SDV rats were made only on the first 30-minutes of the sessions.

**Supplemental Table 1.** Comparisons between pairs of oxytocin doses within each group, on METH infusions. * *p* < 0.05.

| **Sex** | **Surgery** | **0 vs 0.3**  **mg/kg** | **0 vs 1**  **mg/kg** | **0.3 vs 1**  **mg/kg** |
| --- | --- | --- | --- | --- |
| Male | Sham | 0.001* | 0.000* | 0.746 |
|  | SDV | 0.147 | 0.001* | 0.027* |
| Female | Sham | 0.000* | 0.000* | 0.173 |
|  | SDV | 0.135 | 0.004* | 0.010* |

**Supplemental Table 2.** Comparisons between surgery groups at each oxytocin dose on METH infusions. * *p* < 0.05.

| **Sex** | **Surgery** | **Oxytocin dose (mg/kg)** | | |
| --- | --- | --- | --- | --- |
|  |  | **0** | **0.3** | **1** |
| Male | Sham vs. SDV | 0.775 | 0.012* | 0.187 |
| Female | Sham vs. SDV | 0.351 | 0.028* | 0.214 |

**Supplemental Table 3.** Comparisons between METH infusions on each test day and the respective day prior. * *p* < 0.05.

| **Sex** | **Surgery** | **Oxytocin dose (mg/kg)** | | |
| --- | --- | --- | --- | --- |
|  |  | **0** | **0.3** | **1** |
| Male | Sham | 0.475 | 0.001* | 0.003* |
|  | SDV | 0.691 | 0.338 | 0.011* |
| Female | Sham | 0.110 | 0.002* | 0.000* |
|  | SDV | 0.696 | 0.260 | 0.002* |

**Supplemental Table 4.** Comparisons between pairs of oxytocin doses within each group, on METH infusions normalised to a percentage of the day prior. * *p* < 0.05.

| **Sex** | **Surgery** | **0 vs 0.3**  **mg/kg** | **0 vs 1**  **mg/kg** | **0.3 vs 1**  **mg/kg** |
| --- | --- | --- | --- | --- |
| Male | Sham | 0.000* | 0.000* | 0.868 |
|  | SDV | 0.423 | 0.017* | 0.107 |
| Female | Sham | 0.003* | 0.000* | 0.122 |
|  | SDV | 0.192 | 0.000* | 0.000* |

**Supplemental Table 5.** Comparisons between surgery groups at each oxytocin dose on METH infusions normalised to a percentage of the day prior. * *p* < 0.05.

| **Sex** | **Surgery** | **Oxytocin dose (mg/kg)** | | |
| --- | --- | --- | --- | --- |
|  |  | **0** | **0.3** | **1** |
| Male | Sham vs. SDV | 0.784 | 0.005* | 0.114 |
| Female | Sham vs. SDV | 0.273 | 0.016* | 0.101 |

**Supplemental Table 6.** Comparisons between 0 and 0.3 mg/kg oxytocin within each group, on active nose pokes during cue-induced reinstatement, analysed as raw active nose pokes. * *p* < 0.05.

| **Sex** | **Surgery** | **0 vs 0.3**  **mg/kg** |
| --- | --- | --- |
| Male | Sham | 0.011* |
|  | SDV | 0.211 |
| Female | Sham | 0.011* |
|  | SDV | 0.000* |

**Supplemental Table 7.** Comparisons between surgery groups at each dose of oxytocin on cue-induced reinstatement, either as raw active nose pokes, or as active nose pokes following 0.3 mg/kg oxytocin normalised to a percentage of active nose pokes made following 0 mg/kg oxytocin. * *p* < 0.05.

| **Sex** | **Surgery** | **Oxytocin dose (mg/kg)** | | |
| --- | --- | --- | --- | --- |
|  |  | **0** | **0.3** | **0.3 as percentage of 0** |
| Male | Sham vs. SDV | 0.877 | 0.159 | 0.041* |
| Female | Sham vs. SDV | 0.198 | 0.260 | 0.388 |

**Supplemental Table 8.** Comparisons between 0 and 0.3 mg/kg oxytocin within each group, on locomotor activity during cue-induced reinstatement, analysed as locomotor counts following 0.3 mg/kg oxytocin normalised to a percentage of locomotor counts made following 0 mg/kg oxytocin. * *p* < 0.05.

| **Sex** | **Surgery** | **0 vs 0.3**  **mg/kg** |
| --- | --- | --- |
| Male | Sham | 0.429 |
|  | SDV | 0.603 |
| Female | Sham | 0.076 |
|  | SDV | 0.011* |

**Supplemental Table 9.** Comparisons between surgery groups at each dose of oxytocin at cue-induced reinstatement, either as locomotor counts, or as locomotor counts following 0.3 mg/kg oxytocin normalised to a percentage of locomotor counts made following 0 mg/kg oxytocin. * *p* < 0.05.

| **Sex** | **Surgery** | **Oxytocin dose (mg/kg)** | | |
| --- | --- | --- | --- | --- |
|  |  | **0** | **0.3** | **0.3 as percentage of 0** |
| Male | Sham vs. SDV | 0.526 | 0.649 | 0.270 |
| Female | Sham vs. SDV | 0.608 | 0.612 | 0.718 |

**Supplemental Table 10.** Comparison of active nose pokes between pairs of doses, or active nose pokes following 0.3 or 1 mg/kg oxytocin normalised to a percentage of active nose pokes made following 0 mg/kg oxytocin, within each surgery group and sex, during METH-primed reinstatement * *p* < 0.05.

| **Sex** | **Surgery** | **0 vs 0.3**  **mg/kg** | **0 vs 1**  **mg/kg** | **0.3 vs 1**  **mg/kg** | **Percentage of 0 mg/kg:**  **0.3 vs 1 mg/kg** |
| --- | --- | --- | --- | --- | --- |
| Male | Sham | 0.022* | 0.003* | 0.186 | 0.401 |
|  | SDV | 0.704 | 0.169 | 0.781 | 0.314 |
| Female | Sham | 0.059 | 0.002* | 0.036* | 0.026* |
|  | SDV | 0.003* | 0.012* | 0.534 | 0.668 |

**Supplemental Table 11.** Comparison of active nose pokes between pairs of doses, or as active nose pokes following 0.3 or 1 mg/kg oxytocin normalised to a percentage of active nose pokes made following 0 mg/kg oxytocin, between surgery groups for each sex, during METH-primed reinstatement * *p* < 0.05.

| **Sex** | **Surgery** | **Oxytocin dose (mg/kg)** | | | **Percentage of 0 mg/kg** | |
| --- | --- | --- | --- | --- | --- | --- |
|  |  | **0** | **0.3** | **1** | **0.3 mg/kg %** | **1 mg/kg %** |
| Male | Sham vs. SDV | 0.146 | 0.661 | 0.173 | 0.039* | 0.047* |
| Female | Sham vs. SDV | 0.656 | 0.201 | 0.461 | 0.264 | 0.315 |

**Supplemental Table 12.** Comparison of locomotor activity induced by METH-primed reinstatement, or as locomotor activity following 0.3 or 1 mg/kg oxytocin normalised to a percentage of locomotor activity following 0 mg/kg oxytocin between pairs of oxytocin doses, within each surgery group and sex. * *p* < 0.05.

| **Sex** | **Surgery** | **0 vs 0.3**  **mg/kg** | **0 vs 1**  **mg/kg** | **0.3 vs 1**  **mg/kg** | **Percentage of 0 mg/kg:**  **0.3 vs 1 mg/kg** |
| --- | --- | --- | --- | --- | --- |
| Male | Sham | 0.223 | 0.005 | 0.032* | 0.027* |
|  | SDV | 0.098 | 0.003* | 0.297 | 0.268 |
| Female | Sham | 0.004* | 0.000* | 0.010* | 0.004* |
|  | SDV | 0.050* | 0.008* | 0.324 | 0.483 |

**Supplemental Table 13.** Comparison of locomotor activity induced by METH-primed reinstatement, or as locomotor activity following 0.3 or 1 mg/kg oxytocin normalised to a percentage of locomotor activity made following 0 mg/kg oxytocin, between surgery groups for each sex, * *p* < 0.05.

| **Sex** | **Surgery** | **Oxytocin dose (mg/kg)** | | | **Percentage of 0 mg/kg** | |
| --- | --- | --- | --- | --- | --- | --- |
|  |  | **0** | **0.3** | **1** | **0.3 mg/kg %** | **1 mg/kg %** |
| Male | Sham vs. SDV | 0.778 | 0.961 | 0.233 | 0.777 | 0.406 |
| Female | Sham vs. SDV | 0.789 | 0.874 | 0.083 | 0.455 | 0.017* |
